# Stability of Nitroxide Biradical TOTAPOL in Biological Samples

**DOI:** 10.1101/592766

**Authors:** Kelsey McCoy, Rivkah Rogawski, Olivia Stovicek, Ann McDermott

## Abstract

We characterize chemical reduction of a nitroxide biradical, TOTAPOL, used in dynamic nuclear polarization (DNP) experiments, specifically probing the stability in whole-cell pellets and lysates, and present a few strategies to stabilize the biradicals for DNP studies. DNP solid-state NMR experiments use paramagnetic species such as nitroxide biradicals to dramatically increase NMR signals. Although there is considerable excitement about using nitroxide-based DNP for detecting the NMR spectra of proteins in whole cells, nitroxide radicals are reduced in minutes in bacterial cell pellets, which we confirm and quantify here. We show that addition of the covalent cysteine blocker N-ethylmaleimide to whole cells significantly slows the rate of reduction, suggesting that cysteine thiol radicals are important to *in vivo* radical reduction. The use of cell lysates rather than whole cells also slows TOTAPOL reduction, which suggests a possible role for the periplasm and oxidative phosphorylation metabolites in radical degradation. Reduced TOTAPOL in lysates can also be efficiently reoxidized with potassium ferricyanide. These results point to a practical and robust set of strategies for DNP of cellular preparations.

## Introduction

Dynamic nuclear polarization (DNP) solid-state NMR enhances the sensitivity of NMR experiments by polarization transfer from the electron spins of stable free radicals to nearby nuclear spins^1^. Samples of interest, with appropriate isotopic enrichment, are typically mixed with the radical polarizing agent (often a nitroxide), cooled to cryogenic temperatures, and irradiated at the electron Larmor frequency to excite the electronic transitions, which leads to increased nuclear signal. Although a number of polarization transfer mechanisms exist, the cross effect, which couples two electrons to one nucleus^2^, is particularly useful for biological applications at high field. In order to maximize the percentage of spin pairs satisfying the cross effect matching condition, nitroxide biradicals composed of two tethered paramagnetic centers such as TOTAPOL^3^ and AMUPOL^4^ have been developed (see Figure 1). These biradicals have been used to enhance the NMR signals of a wide range of biological samples available in limited quantities, such as the DNA component of the Pf1 bacteriophage^5^ and microgram quantities of human GPCR membrane proteins^6,7^, yielding insight into the conformations of dilute components on a practical timescale.

**Figure 1:**
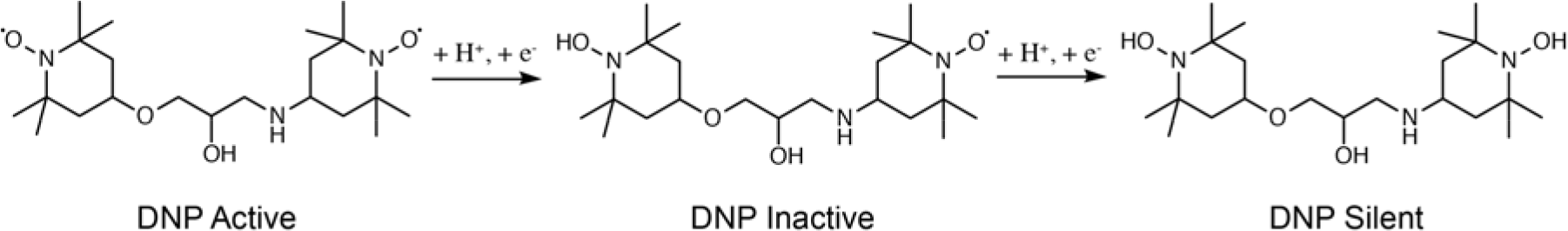
Chemical structure of paramagnetic TOTAPOL^3^ (left) and illustration of its reduction to the DNP inactive (providing DNP enhancement only if in close proximity to another radical) monoradical and DNP silent (incapable of providing DNP enhancement) hydroxylamine forms.

An exciting application of this sensitivity enhancement is its use in in-cell NMR experiments that directly probe protein conformations and interactions in native cellular environments. A number of intriguing results suggest that the signal enhancement provided by DNP can enable studies of proteins in whole cells or lysates^8–11^. However, a challenge for these experiments is the fast reduction of free radicals in cellular environments. This challenge has been well documented by EPR^12–15^, where stable free radicals are used for in-cell investigation of protein conformations. For example, Summerer and coworkers showed that the spin labeled unnatural amino acid SLK-1 was reduced in both whole *E. coli* cells and lysates^16^. When Cafiso *et al.* labeled whole *E. coli* cells with the nitroxide spin label MTSSL, they observed no signal for cysteine mutants located in the periplasm, implying that reduction was fast upon uptake^17,18^. Comparison of reduction rates for different nitroxide compounds indicates that piperidine nitroxides, the building block of most DNP polarizing agents, can be reduced on the order of minutes^13^. For example, the lifetime of the piperidine nitroxide TEMPOL in *Xenopus laevis* oocyte cell extracts is only 3.8 minutes^12^.

Studies of the mechanism of nitroxide reduction in biological samples indicate that sulfhydryl groups on proteins and glutathione play a larger role in reduction than endogenous ascorbic acid^1,19^, although radical oxygen species can also contribute to nitroxide signal decay^19^. Treatment with alkaline solutions^12^, pre-treatment with cysteine specific blockers^20^, and pre-treatment with protease inhibitors^15^ have all been used to protect against nitroxide reduction. It is also possible to reoxidize reduced nitroxides using potassium ferricyanide^21,22^.

Due to the reliance of cross-effect DNP on stable biradical pairs, reduction of the biradicals by the cell quickly leads to failure of the cross-effect DNP experiment. Even partial reduction of the nitroxide moiety to form the corresponding hydroxylamine leads to a loss of DNP enhancements. For example, 50% random reduction of nitroxide centers will lead to a 75% loss of *biradical centers*, i.e. creation of compounds with at least one member of the biradical reduced (Figure 1). In fact, native membranes suspended with TOTAPOL have shown loss of DNP enhancement over time^23^, possibly due to biradical reduction by free cysteines.

To successfully carry out DNP experiments on whole cells or cell lysates, the nitroxide must remain paramagnetic for at least the time it takes to mix it thoroughly with the sample, insert the sample into a rotor, and freeze. This can be on the order of 10 minutes to 1 hour, depending on ease of sample packing and success of magic angle spinning. As mentioned above, under some circumstances piperidine nitroxides can be reduced on the order of minutes. Therefore, it is important both to quantify the timescale of nitroxide biradical reduction in biological samples and to find strategies to combat reduction. Here we investigate the lifetime of the DNP-relevant nitroxide biradical TOTAPOL in *E. coli* cell pellets, suspensions, and lysates. We also test the efficacy of strategies to prolong the lifetime of radicals in cell pellets and lysates, including incubation with N-ethylmaleimide and treatment with potassium ferricyanide.

## Materials and Methods

### Preparation of Whole-Cell Pellets and Lysates for EPR

BL21(DE3) cells transformed with the pUC19 plasmid were inoculated into 100 mL LB cultures from an overnight pre-culture and grown to an OD_600_ of ~0.8. Cells were collected by centrifugation at 3,000 × g for 10 minutes, re-suspended in 10 mL PBS, and incubated at 4 °C for 30 minutes. Approximately 400 mg of cells were collected from 100 mL of cell culture. For experiments with N-ethylmaleimide (NEM), cells were incubated for 30 minutes at 4 °C in 10 mL PBS containing NEM. Cells were then pelleted by centrifugation, re-suspended in 1 mL PBS, aliquoted into individual Eppendorf tubes, and pelleted by centrifugation in a benchtop microcentrifuge at 11,700 × g. The supernatant was removed, and individual cell pellets were kept on ice until the TOTAPOL was added. The approximate mass of each cell pellet was 30 mg. See Figure S1 for detailed workflow.

For lysate experiments, the pellet from 100 mL of cell culture grown as described above was re-suspended in 1 mL lysis buffer (50 mM Tris/HCl pH 8, 3.22 mM EDTA). Cells were incubated on ice for 30 minutes and then lysed with sonication (5 rounds of 10 s sonication alternating with a 90 s rest on ice). This preparation was used for crude lysate experiments. For clarified lysate experiments, the lysate was spun down in a benchtop microcentrifuge at 11,700 × g, and the supernatant was used for the experiment. For the whole cell suspension experiment, cells were simply re-suspended in 1 mL PBS and incubated on ice for 30 minutes, and this suspension was used without further treatment.

TOTAPOL was added to the samples by gently mixing in the appropriate amount of a 50 mM TOTAPOL solution in DMSO with a pipette tip. At each time point (~5 min, 10 min, 15 min, 30 min, 45 min) a *separate* pellet or aliquot was diluted to a final volume of 1 mL in PBS, bringing the final TOTAPOL concentration down to the micromolar regime, and the spectrum was collected. For potassium ferricyanide experiments, the appropriate amount of a 1M K_3_(Fe(CN)_6_) dissolved water was added to the sample being measured approximately two minutes prior to measuring the spectrum.

### EPR Experiments

CW EPR data was acquired at room temperature using a Bruker EMX X-band EPR spectrometer at a microwave frequency of 9.756 GHz, with a center field of 3480 G and sweep width of 100 G. A modulation frequency of 100 kHz, modulation amplitude of 1.00 G, conversion time of 164 seconds, and microwave power of 20.1 mW were used. These parameters were optimized on a solution of TOTAPOL to ensure that lineshape and intensity did not vary with parameter choice, and conditions chosen to maximize S/N.

For each sample, the doubly integrated EPR spectral intensity was normalized to the integrated intensity of a control TOTAPOL sample in PBS. A new unreduced TOTAPOL sample was prepared fresh for each experiment to account for instrument instability and variation in TOTAPOL stock solutions.

## Results and Discussion

### TOTAPOL Is Reduced in Cell Pellets in Minutes

Initial DNP experiments on cell pellets with TOTAPOL yielded no DNP signal enhancement (data not shown), despite the addition of biradical stocks known to be paramagnetic when tested with EPR. We hypothesized that this was due to nitroxide reduction specifically by the cellular environment, since nitroxide radicals, the building blocks of the TOTAPOL DNP polarizing agent, are known to be stable in solution. We confirmed this stability by tracking the intensity of a 0.1 mM TEMPO solution over the course of a month, and found that well over 90% of the overall integrated intensity was maintained even when stored at room temperature (Figure S2). We therefore concluded that most of the observed TOTAPOL reduction was caused by reducing agents in the cellular environment.

We proceeded to use EPR to track the stability of DNP-relevant concentrations of TOTAPOL in whole cells, with the aim of manipulating the timescale of reduction to facilitate the preparation of whole-cell DNP samples. We designed an experiment to simulate the conditions the radical experiences inside a DNP rotor packed with whole cells, where small amounts of radical are mixed with a tightly packed cell pellet and incubated at room temperature for a variable amount of time (see Materials and Methods). CW X-band EPR spectroscopy was used to measure the amount of paramagnetic radical left by calculating the total integrated intensity of the spectrum^12^.

Our reduction curves do not fit well to monoexponential decay curves (Figure S3), particularly in cases when the signal intensity does not approach zero at the end of the sampling period. This is not unexpected for a bimolecular reaction such as radical reduction and is consistent with the non-exponential decay monoradical nitroxides in cell extracts previous reported^24^. Moreover, it is unlikely that any single reducing agent is responsible for the majority of TOTAPOL reduction in either cell pellet or lysate preparations. The radicals are likely reduced by a complex mixture of small molecules such as glutathione and ascorbate, redox mediators such as quinones, and redox-active enzymes^25^. Therefore, instead of fitting to an exponential decay, we approximated the initial decay rate by fitting a linear regression line to the first three time points (unreduced stock solution, 5 minute incubation, and 10 minute incubation) of each curve (Figure 2; Table S1). This method also allowed us to focus solely on the radical reduction that occurs on the timescale of DNP sample preparation.

**Figure 2:**
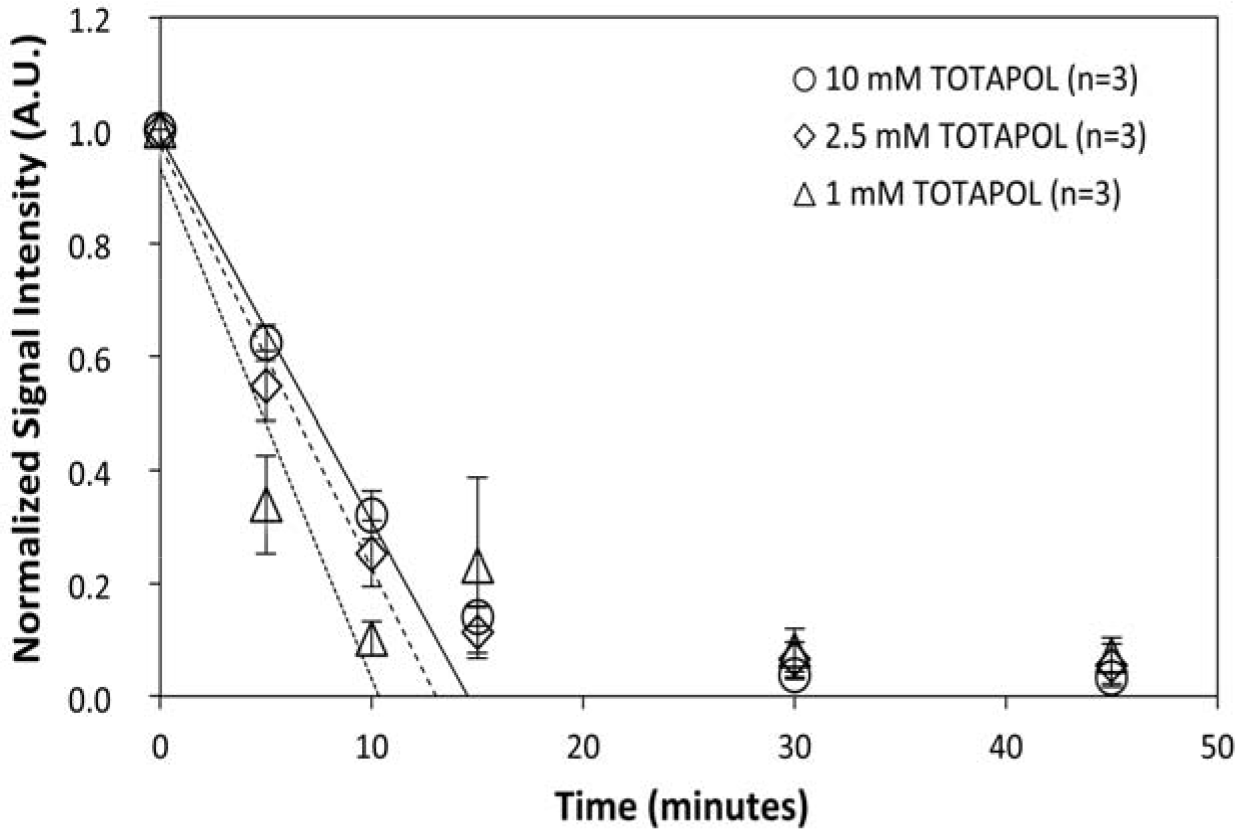
Time course of TOTAPOL reduction in whole-cell pellets for 10 (solid fit line), 2.5 (dashed fit line), and 1 mM (dotted fit line) TOTAPOL. All reduction values are reported as the integrated EPR signal intensity normalized to a standard sample of TOTAPOL. Error bars are plotted as standard error. Best-fit line represents simple linear regression through the first three points (unreduced TOTAPOL, 5 min, 10 min), demonstrating how decay rates are calculated.

At room temperature, whole cells significantly reduce TOTAPOL within minutes. After 10 minutes, only 25 ± 6% of the initial TOTAPOL signal remained. Assuming that radical reduction is random, this corresponds to a biradical concentration of 6.4 ± 3.0 % after 10 minutes (Figures 2,3), and represents a decay rate of 0.18 ± 0.01 mmol TOTAPOL/(L*min) over the first 10 minutes (Table S1). These results demonstrate that within minutes of exposure to densely packed whole cells, the TOTAPOL biradical is significantly reduced, rendering the vast majority of the TOTAPOL DNP-inactive.

**Figure 3:**
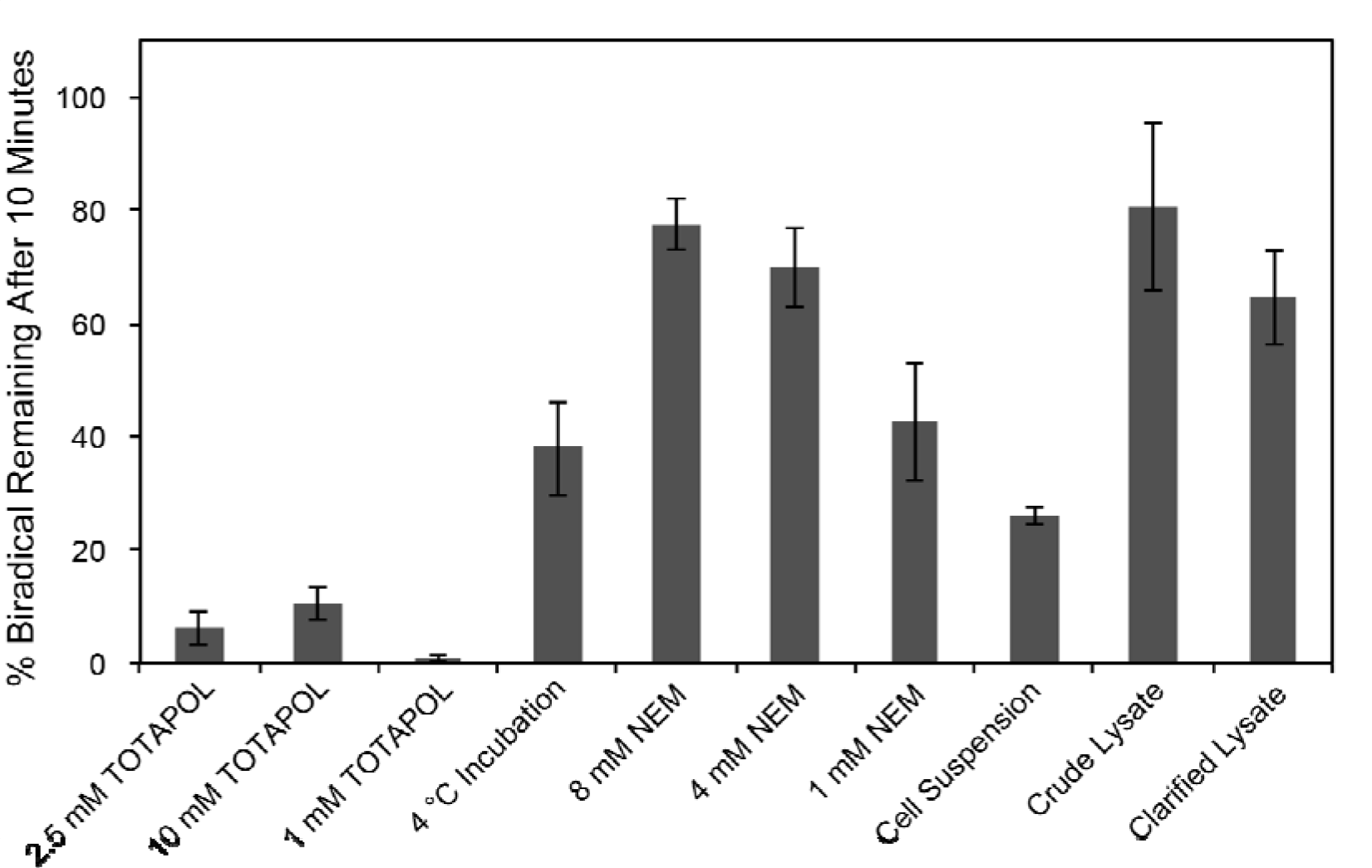
Graph of percentage active biradical remaining after ten minutes for each experimental condition. Cells and lysates were incubated at room temperature with 2.5 mM TOTAPOL unless otherwise indicated. N = 3 for all conditions. Percentage biradical was calculated from the percentage EPR signal remaining assuming random reduction of each radical center. Error plotted as standard error.

Changing the concentration of TOTAPOL caused a proportional change in the reduction rate, indicating first-order reaction kinetics with respect to TOTAPOL (Table S1; Figure S4). This result is in contrast to that observed by Drescher and coworkers^12^, who observed full saturation of the reaction rate for the pyrollidine nitroxide PCA and partial saturation for the piperidine TCA in *Xenopus leavus* cell extract. Our data do not suggest the enzymatic reduction pathway that they propose, but rather agree with previous characterization of cellular reduction from glutathione and other small molecules, a claim supported by our results with N-ethylmaleimide (see below).

### TOTAPOL Reduction Is Slowed in Lysates

Cellular lysates and extracts have been used to approximate the cellular environment for DNP experiments^8, 26^.We therefore tested the effect of cell lysis on the TOTAPOL reduction rate. To control for dilution effects in the lysate, we prepared whole-cell suspensions at the same concentration as the cell lysates. As expected, the suspensions do not reduce TOTAPOL as fast as the cell pellets do (0.12 ± 0.01 mmol/(L*min) versus 0.18 ± 0.01 mmol/(L*min)). This difference can be attributed to the dilution of the cells relative to the TOTAPOL in the suspensions.

However, both crude and clarified lysates (lysates with and without the membrane components, respectively; see Materials and Methods) reduce the TOTAPOL significantly slower than the whole-cell suspensions at all time points (0.043 ± 0.008 mmol/(L*min) and 0.043 ± 0.004 mmol/(L*min) respectively). This is likely due to the destruction of the redox gradients in the periplasm that drive the electron transport chain^26^, and because of the disruption of crowding effects that keep enzymes responsible for turning over reducing agents in close proximity to oxidized reducing agents. Interestingly, both crude and clarified lysates had similar initial TOTAPOL decay rates, suggesting that the presence of the membrane and membrane-associated factors in the crude lysate is less important than the way the reducing equivalents are concentrated and maintained within the cell. These results collectively indicate that radical reduction is not as significant in lysate preparations as it is in whole cells.

We noted, however, that even in clarified lysates, only 64 ± 8 % of biradical centers from a starting concentration of 2.5 mM TOTAPOL remain after 10 minutes at room temperature. While using a higher starting concentration of TOTAPOL does increase the moles of paramagnetic radical left in the pellet, this strategy is not effective for targeted DNP, where stoichiometric concentrations of affinity reagent and target biomolecule are used to avoid enhancement of background signals. It is unlikely that reduction of the TOTAPOL moiety will affect binding of the probe to the target, and therefore a large percentage of reduced probe will lead to significant losses in enhanced signal intensity, since many target biomolecules will be bound to DNP-inactive polarizing agents, and low starting concentrations of radical make it unlikely that another radical will be in close enough proximity to polarize the target biomolecule. Motivated by the desire to enable DNP at a broad range of biradical concentrations, and particularly in the low millimolar to micromolar range, we explored various methods to prevent TOTAPOL reduction in whole cells and lysates.

### TOTAPOL Reduction Is Slowed at 4 °C

We first looked at the effect of lowering the temperature on TOTAPOL reduction in cells. We expected that a lowered temperature would confer a protective effect on the radicals, as both radical reductions by the extant reducing agents within the cell as well as the metabolic processes that maintain the redox balance are slower at near-freezing temperatures. As expected, keeping the cells at 4 °C during incubation with the radical slows down reduction. The initial decay rate slowed to 0.12 ± 0.03 mmol/(L*min), with 38 ± 8 % of the biradical remaining after 10 minutes (Figure 3). However, this increase is not sufficient to reliably and reproducibly maintain stable biradicals for cross-effect DNP, particularly with lower starting concentrations of radical.

### Pre-Treatment with N-ethylmaleimide Significantly Prolongs Nitroxide Lifetime

We tested whether pre-treatment of cells with N-ethylmaleimide (NEM), a cysteine-specific blocker that covalently binds to active thiol groups, could neutralize pools of redox-active cysteines and protect against nitroxide reduction^20,28^. Incubating the cells with 8 mM NEM for 30 minutes prior to the addition of TOTAPOL significantly increased the radical lifetime in whole-cell pellets. The decay rate slowed to 0.028 0.016 mmol/(L*min), corresponding to 78 5 % of active biradical remaining after 10 minutes, demonstrating a significant protective effect relative to untreated cells. We also tested the protective effect of lower concentrations of NEM. 4 mM NEM had a similar effect as 8 mM NEM, but 1 mM NEM was much less effective, with only 42 10 % of active biradical remaining after 10 minutes.

We also tested the effect of NEM treatment on cell lysates. Importantly, for clarified lysates prepared from cells pre-treated with NEM, we observed no measurable reduction over 10 minutes and minimal reduction over 45 minutes (Figure S6). This suggests that cell lysates pre-treated with NEM can be reliably used to preserve biradicals, even when relatively minimal reduction is undesirable, such as when low starting concentrations of TOTAPOL are employed.

### Potassium Ferricyanide Effectively Reoxidizes TOTAPOL in Lysates

One technique for combating the reduction of nitroxide spin labels in whole-cell EPR experiments is to reoxidize reduced radicals with potassium ferricyanide immediately prior to collecting EPR spectra^29,30^. We investigated whether this method could be used to reoxidize reduced TOTAPOL by adding 4- and 20-fold molar excesses of potassium ferricyanide to whole-cell and lysate samples incubated with TOTAPOL for a range of times (see Materials and Methods).

We found that in whole-cell pellets, using a 4-fold molar excess of ferricyanide led to a measurable increase in the detectable TOTAPOL signal intensity at the 45 minute time point but not at the 10 minute time point (Figure 4; Table S2). Increasing the ratio to a 20-fold molar excess of ferricyanide to TOTAPOL led to a further increase, with 50% of the initial TOTAPOL signal observed after a 10-minute incubation followed by ferricyanide reoxidation. However, this 50% reoxidation was the maximum observed in the whole-cell pellets.

**Figure 4:**
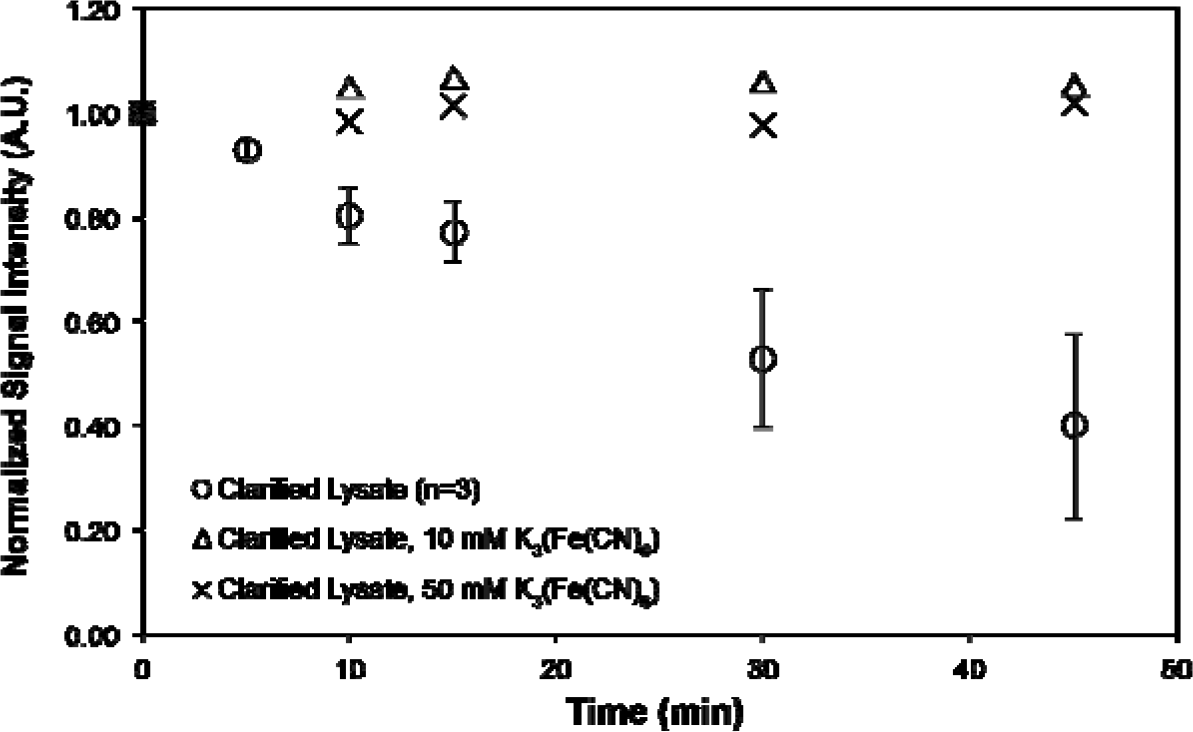
Time course of TOTAPOL reduction in clarified lysates (n = 3; error plotted as standard error) and lysates treated with 10 mM and 50 mM K_3_(Fe(CN)_6_) approximately 2 minutes prior to measurement (n = 1), demonstrating that reduced TOTAPOL can be completely reoxidized in lysates.

However, in lysates, both 4-fold and 20-fold molar excesses of potassium ferricyanide led to complete or near-complete reoxidation of the reduced TOTAPOL at all time points (Figure 4; Table S2). This effect was observed in both crude and clarified lysates, demonstrating that TOTAPOL can be reliably reoxidized in a variety of lysate preparations. This provides an alternative strategy for preparing DNP samples in lysates, if the covalent cysteine chemistry of NEM treatment is not desired for a given experiment or longer sample preparation times are necessary. Our data support previous work by Saxena and coworkers that found that the nitroxide monoradical PCA’s reduction by *Xenopus laevis* oocyte extracts was slowed by the addition of up to 5 molar equivalents K_3_(Fe(CN)_6_) ^30^, along with their recent result that reduced TEMPOL in *X. laevis* oocytes can be reoxidized by injection of K_3_(Fe(CN)_6_)^22^.

Previous work by the Hediger group showed that in the Gram-positive bacteria *Bacillus subtilis*, TOTAPOL preferentially binds to the peptidoglycan cell wall.^31^ Ferricyanide is known to be taken up into *E. coli* cells and interacts with the electron transport chain and is reduced in the periplasmic space.^32,33^ Taken together with our results that ferricyanide efficiently reoxidizes TOTAPOL in lysates but only partially reoxidizes TOTAPOL in whole cells, and that it also reoxidizes TEMPOL when co-injected into *X. laevis*^30^ oocytes, this suggests that TOTAPOL is being reduced primarily in the periplasmic space, rather than in the cytoplasm.

### NEM Increases TOTAPOL Concentration in DNP Samples

We prepared whole cell DNP samples using our method of NEM treatment and measured the EPR spectra at the time rotor packing. We found that samples washed with 8 mM NEM had ~2× as much TOTAPOL at the time of sample freezing than untreated samples, across multiple conditions (Table S3). Surprisingly, this did not correlate to increased DNP enhancements, which we believe is due to poor TOTAPOL uptake, or preferential uptake of TOTAPOL into the periplasmic space (see supplementary materials). Biradical uptake remains a challenge, and further work is needed in this area before true in cell DNP can be achieved.

## Conclusions

DNP provides a powerful tool for extending solid-state NMR studies into native cellular environments, but requires stable and biocompatible radicals. Nitroxides are popular, but are quickly reduced in typical cellular environments. We have characterized the timescale of TOTAPOL reduction in whole cells and in lysates and demonstrate that it is fast relative to the typical DNP sample preparation protocol timescale. We present a number of strategies for combating reduction, including treatment of cells with covalent cysteine blocker N-ethylmaleimide and reoxidation of reduced radicals with potassium ferricyanide in lysates. Using these strategies, we have demonstrated that TOTAPOL’s lifetime can be extended in the context of whole cell DNP samples.

## Acknowledgements

This work was supported by a grant from the National Science Foundation (NSF): MCB1412253 to A.E.M and by the National Institute of Health Grant P41GM118302 for the Center on Macromolecular Dynamics by NMR Spectroscopy located at the New York Structural Biology Center (NYSBC). A.E.M is a member of the NYSBC, and the data collected at NYSBC was enabled by a grant from NYSTAR and ORIP/NIH facility improvement grant CO6RR015495. The 600 MHz DNP/NMR spectrometer was purchased with funds from NIH grant S10RR029249. R.R was supported by the National Institute of Health Training Program in Molecular Biophysics T32GM008281. K.M.M was supported by a NSF Graduate Fellowship DGE 16-44869.

The authors thank Dr. Steffen Jokusch for help and use of the CW X-band EPR instrument, and Drs. Boris Itin and Mike Goger of the NYSBC for help with DNP instrumentation.

---

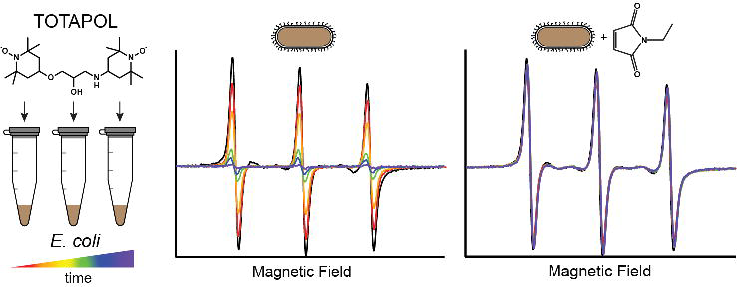

