## Supplementary Information for "Stability of Nitroxide Biradical TOTAPOL in Biological Samples"

**Supplemental Methods:**

*DNP Experiments*

For DNP experiments, 100 mL of perdeuterated cell culture ($OD_{600}$ >2.0) over expression either *E. coli* DHFR or *E. coli* TIM was harvested by centrifugation at 4,150 x g in a Sorvall benchtop centrifuge. Approximately 400 mg of cells were collected. The cells were rinsed twice with a 5 mL aliquot of PBS, with and without 8 mM NEM. Cell pellets were then collected in Eppendorf tubes. Cell pellets of approximately 30 mg were mixed with a combination of a 50 mM stock solution of TOTAPOL in DMSO and buffer such that the final TOTAPOL concentration was 10 mM, 5 mM, or 1 mM (10% final DMSO concentration in all cases). The cell pellets were incubated at room temperature for 10 minutes prior to pelleting into the rotor in order to test the efficacy of NEM treatment. The cell pellet was collected by centrifuging directly into a 3.2 mm sapphire rotor and flash frozen in liquid nitrogen.

DNP experiments were performed on a 600 MHz (14.1 T) Bruker Avance III-DNP system at the New York Structural Biology Center (NYSBC), equipped with a 395 GHz gyrotron and an HCN triple channel E-free probe. Experimental parameters were as follows: MAS rates of 11 kHz, LTMAS temperatures of 100.3 K for variable temperature gas, 105.6 K for bearing gas, 102.9 K for drive gas, and 110.0 K for the VT temperature sensor.

*Nitroxide Uptake*

TOTAPOL uptake into BL21(DE3) E. coli cells was assayed using paramagnetic broadening reagent chromium (II) oxalate (CrOx). CrOx is excluded from cells, and thus broadens only the TOTAPOL lines that originate from outside of the cells, allowing uptake to be assayed using CW EPR (Samuni et al. 1991). The presence of peaks in the spectrum after addition of CrOx indicates uptake.

Uptake was measured by preparing protonated BL21(DE3) cells overexpressing TIM in the same manner as for preparation of DNP samples. Cells were incubated with a final concentration of 10 mM TOTAPOL (at a final concentration fo 10% DMSO) at room temperature for 10 minutes and then brought to 200 mM CrOx immediately before measuring the CW EPR spectrum.

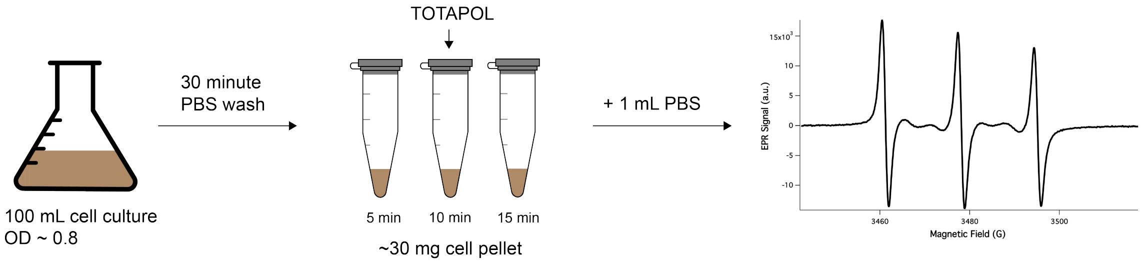

**Figure S1**: Experimental workflow for whole cell pellet EPR experiments. Cells were pelleted, washed with PBS buffer, pelleted again, and TOTAPOL was added. Note that separate samples were prepared for each time point.

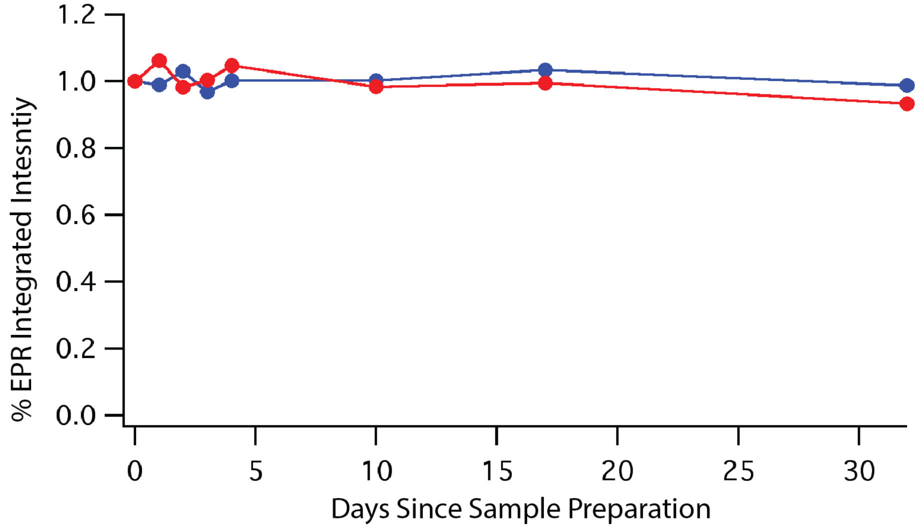


**Figure S2:** Plot of the integrated signal intensity of a 100 μM solution of TEMPO in PBS at pH 7.4 over 30 days indicates minimal reduction of the nitroxide moiety in buffered solution. Curve in red represents the decay for an EPR sample stored at room temperature, and the curve in blue represents the decay for an EPR sample stored at 4 °C between EPR measurements, which were conducted at room temperature.

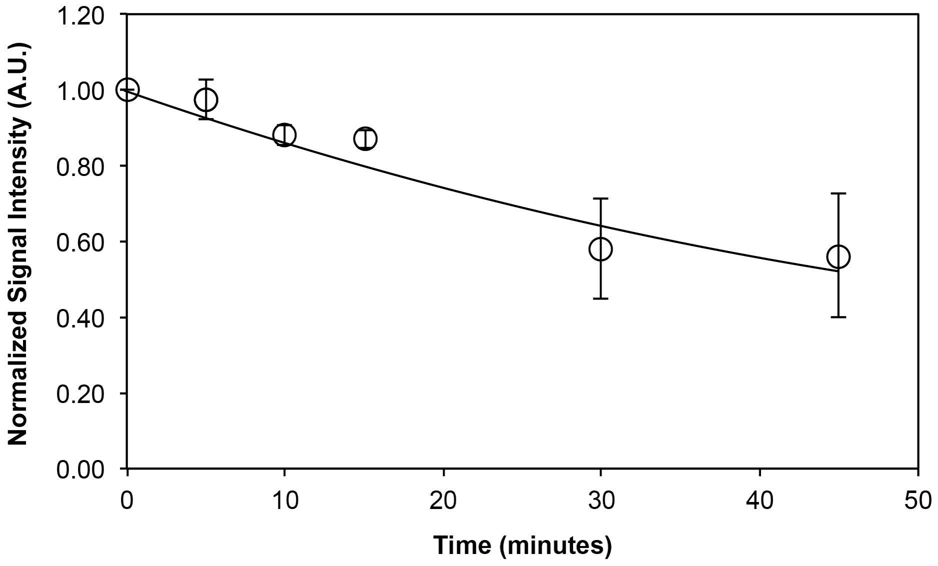


**Figure S3:** Representative decay of normalized integrated EPR intensity of TOTAPOL in a whole cell pellet (shown with the 8 mM NEM dataset) as a function of time does not fit well to a single exponential decay curve in cases where the signal intensity does not approach zero.

| | Sample | Concentration TOTAPOL (mM) | Other Details | Decay Rate (mmol/L*min) | % Biradical Remaining, 10 min |
|---|---|---|---|---|---|
| 1 | Whole cells | 2.5 | | 0.18 ± 0.0080 | 6.4 ± 5.1 |
| 2 | Whole cells | 2.5 | 4 °C | 0.12 ± 0.028 | 38.3 ± 14.4 |
| 3 | Whole cells | 10 | | 0.65 ± 0.019 | 10.3 ± 4.8 |
| 4 | Whole cells | 1.0 | | 0.095 ± 0.0054 | 1.0 ± 1.1 |
| 5 | Whole cells | 2.5 | 8 mM NEM | 0.028 ± 0.016 | 77.6 ± 7.9 |
| 6 | Whole cells | 2.5 | 4 mM NEM | 0.048 ± 0.0048 | 69.9 ± 12.0 |
| 7 | Whole cells | 2.5 | 1 mM NEM | 0.070 ± 0.0088 | 42.4 ± 18.0 |
| 8 | Cell Suspension | 2.5 | | 0.12 ± 0.0068 | 26.0 ± 2.5 |
| 9 | Crude Lysate | 2.5 | | 0.043 ± 0.0083 | 80.9 ± 25.8 |
| 10 | Clarified Lysate | 2.5 | | 0.043 ± 0.0045 | 64.3 ± 14.5 |

**Table S1:** EPR signal decay rates for all samples tested, given in units of mM TOTAPOL/min, and percentage active biradical remaining after 10 minute incubation, calculated assuming random reduction of radical centers (error given as standard deviation). All conditions listed are for whole cell pellets incubated at room temperature unless specified. N = 3 for all conditions.

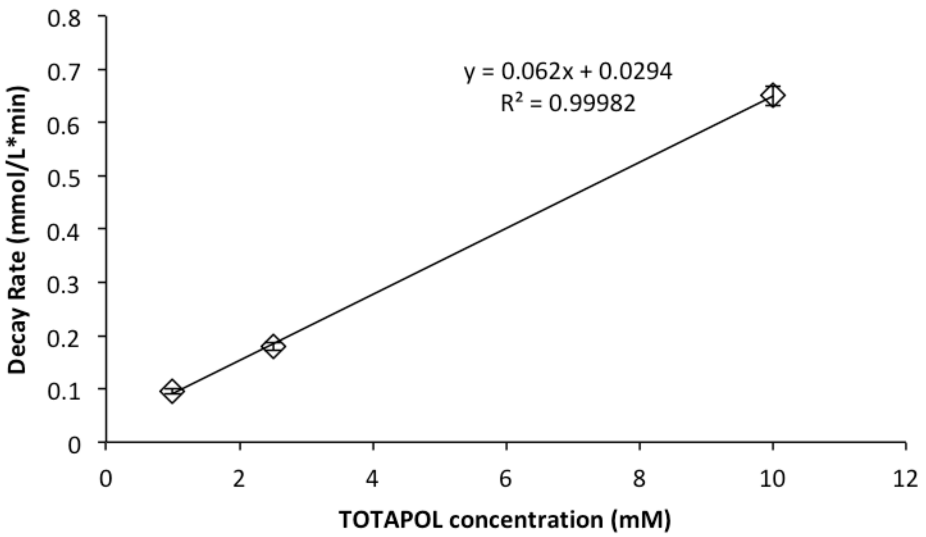


**Figure S4:** Molar decay rate as a function of TOTAPOL concentration is well fit by a line (with an $R^2$ of 0.99), indicating that the decay rate is first order with respect to TOTAPOL concentration for a fixed concentration of cells.

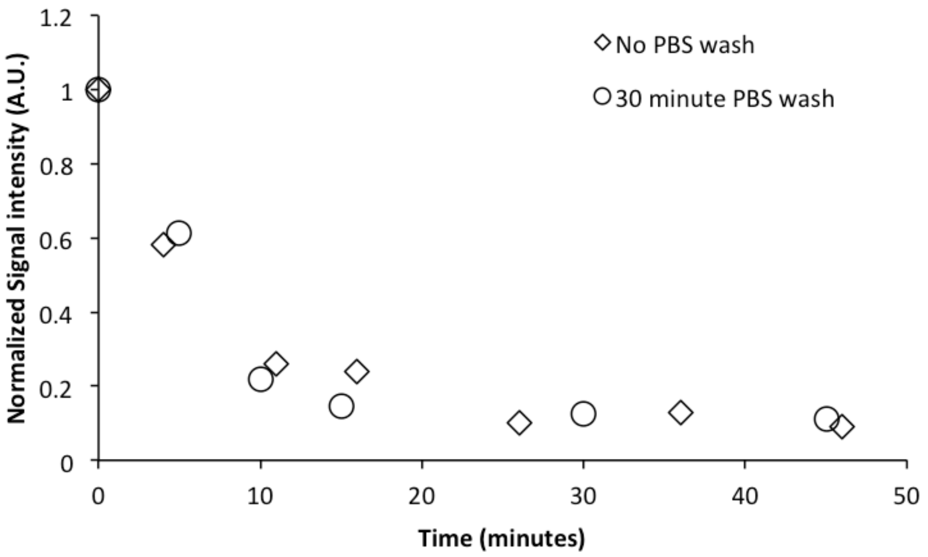


**Figure S5:** 2.5 mM TOTAPOL reduction in whole cell pellets incubated at room temperature with and without a 30-minute incubation in PBS prior to the addition of TOTAPOL indicates that the curve and rates are very similar, i.e. that the PBS wash has a minimal effect on the reducing capability of the cells.

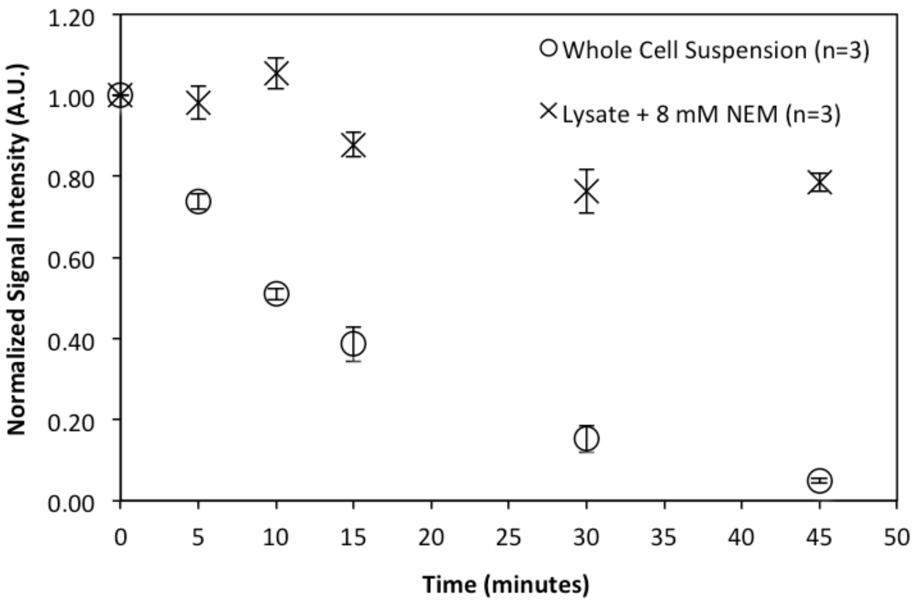


**Figure S6:** 2.5 mM TOTAPOL reduction in lysates pretreated with 8 mM NEM compared to TOTAPOL reduction in whole cell suspensions of similar concentration. No significant reduction was observed after 10 minutes, and minimal reduction was observed at 45 minutes. Error plotted as standard error.

| | Sample | Concentration TOTAPOL (mM) | Concentration $K_3(Fe(CN)_6)$ (mM) | % Signal Remaining, 10 min | % Signal Remaining, 45 min |
|---|---|---|---|---|---|
| 1 | Whole cells | 2.5 | 10 | 18 | 17 |
| 2 | Whole cells | 2.5 | 50 | 52 | 23 |
| 3 | Crude Lysate | 2.5 | 10 | 93 | 120 |
| 4 | Crude Lysate | 2.5 | 50 | 96 | 88 |
| 5 | Clarified Lysate | 2.5 | 10 | 105 | 106 |
| 6 | Clarified Lysate | 2.5 | 50 | 98 | 102 |

**Table S2:** TOTAPOL reoxidation with $K_3(Fe(CN)_6)$ in whole cells and lysates, measured as the percent of initial EPR signal remaining after 10 and 45 minutes of TOTAPOL incubation (compare to Table 1 in the main manuscript). This data indicates that $K_3(Fe(CN_6))$) does reoxidize a population of the radical in the whole cells, and most of the radical that is reduced in the lysate. A two-minute incubation with either 4 or 20 molar excess $K_3(Fe(CN)_6)$ was not sufficient to fully reoxidize the reduced TOTAPOL in whole cell pellets. However, both ferricyanide concentrations were sufficient to reoxidize most of the TOTAPOL in both crude and clarified lysates.

*NEM Does Not Increase DNP Enhancements*

We measured DNP enhancement factors on cell pellets with 10 mM, 5 mM, and 1 mM TOTAPOL prepared with and without pretreatment with NEM (see Supplementary Methods). We incubated the cell pellets with the TOTAPOL for 10 minutes at room temperature prior to stimulate TOTAPOL uptake into the cells and measured the final TOTAPOL concentrations using EPR (Table S3). We did not observe different enhancement factors between the samples treated with NEM and the samples without NEM, despite measuring ~2x as much EPR signal in the samples treated with NEM. DNP enhancement factors were found to be dependent solely on the starting TOTAPOL concentrations, not the final TOTAPOL concentrations in the samples, across all samples tested (Figure S6 and Table S3).

Samples were prepared using two different overexpressed proteins (DHFR in the sample treated with 10 mM TOTAPOL and triosephosphate isomerase (TIM) in the 5 mM and 1 mM TOTAPOL samples), indicating that this effect is not due to the specifics of protein overexpression. In all three cases, the DNP enhancements were within experimental variability with and without NEM treatment, however the enhancements did roughly scale with starting TOTAPOL concentration (i.e. 10 mM TOTAPOL samples had roughly the same enhancements with and without NEM, which were larger than the enhancements in the 5 mM TOTAPOL samples). However, due to the qualitative nature of using DNP enhancements and CW EPR measurements as read outs, it is difficult to say exactly how they relate to each other.

Further investigation into this effect revealed that the 10% DMSO we added to our samples to stimulate uptake were causing protein to leak out of the cells, while at the same time not increasing TOTAPOL uptake (Figure S7). This suggests that the majority of the TOTAPOL reduction we observed is taking place in the media rather than the cells, and that the observed enhancements are from protein that has leaked into the media through holes created by the DMSO. Further investigation must be done into radical uptake into whole cells, however, regardless of radical uptake, we have shown that NEM reliably prevents radical reduction and can be used to preserve biradicals such as TOTAPOL in cellular contexts.

| Sample | Initial [TOTAPOL] | Final [TOTAPOL] | NEM | ε on Co |
|---|---|---|---|---|
| DHFR | 10 mM | 2.2 mM | + | 11 |
| DHFR | 10 mM | 1.1 mM | - | 16 |
| TIM | 5 mM | 1.0 mM | + | 9.5 |
| TIM | 5 mM | 0.57 mM | - | 8 |
| TIM | 1 mM | 0.12 mM | + | 3.3 |
| TIM | 1 mM | 0.045 mM | - | 3.3 |

**Table S3:** TOTAPOL concentrations and DNP enhancements on protein carbonyls in a series of whole cell DNP samples with and without NEM treatment. NEM treatment increased the final TOTAPOL concentration in the samples (verified using EPR) twofold, but DNP enhancements did not increase commensurately. DNP enhancements are weakly correlated to initial TOTAPOL concentration but show no increase upon NEM treatment.

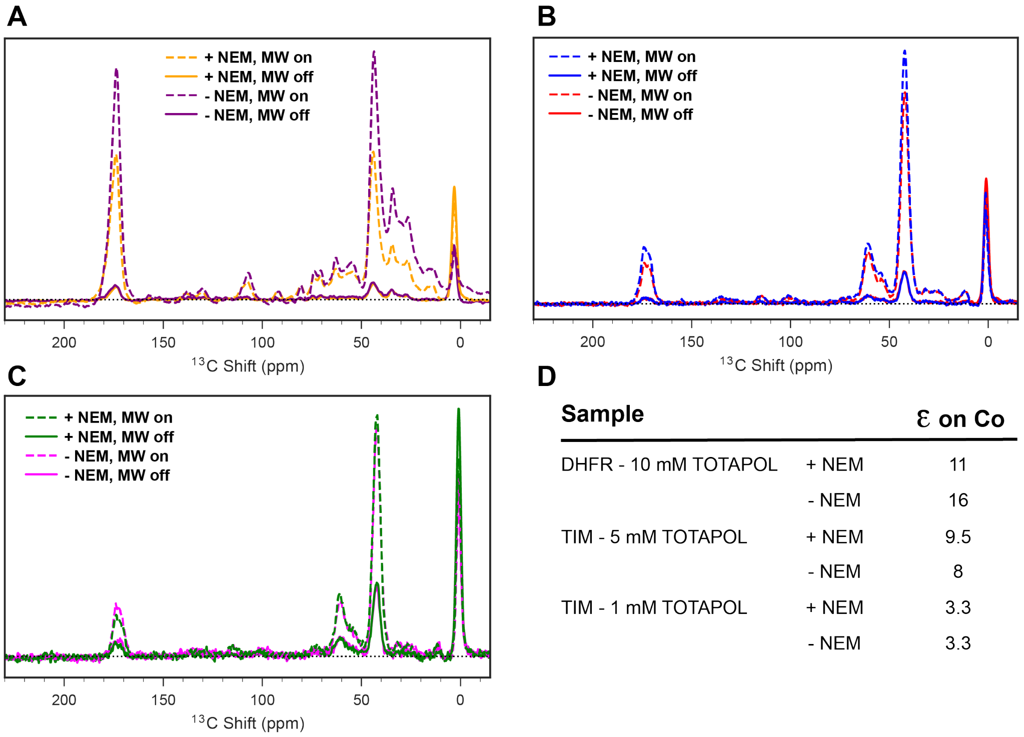


| Sample | | ε on Co |
|---|---|---|
| DHFR - 10 mM TOTAPOL | + NEM | 11 |
| | - NEM | 16 |
| TIM - 5 mM TOTAPOL | + NEM | 9.5 |
| | - NEM | 8 |
| TIM - 1 mM TOTAPOL | + NEM | 3.3 |
| | - NEM | 3.3 |

**Figure S6:** DNP enhancements of protein signals in whole cell pellets is roughly equivalent in samples with and without NEM treatment. A) $^{13}$C CP spectra of BL21(DE3) whole cells expressing $^{13}$Co-Ile, $^{15}$N-Ala, $^{15}$N/$^{13}$Ca-Gly DHFR in a fully deuterated background, with and without NEM pretreatment. 10 mM TOTAPOL was used. Spectra demonstrate enhancements of 11 and 16 respectively B) $^{13}$C CP spectra of BL21(DE3) whole cells expressing $^{13}$Co-Ile, $^{15}$N-Ala, $^{15}$N/$^{13}$Ca-Gly TIM in a fully deuterated background, with and without NEM pretreatment. 5 mM TOTAPOL was used. Spectra demonstrate enhancements of 9.5 and 8 respectively. C) $^{13}$C CP spectra of BL21(DE3) whole cells expressing $^{13}$Co-Ile, $^{15}$N-Ala, $^{15}$N/$^{13}$Ca-Gly TIM in a fully deuterated background, with and without NEM pretreatment. 1 mM TOTAPOL was used. Spectra demonstrate enhancements of 3.3 in both cases. All spectra were collected at 600 MHz $^{1}$H frequency with an MAS frequency of 11 kHz and sample temperature of 110 K, using $^{1}$H-$^{13}$C CP with 512 scans (A) or 128 scans (B,C) and a recycle delay of 3 seconds

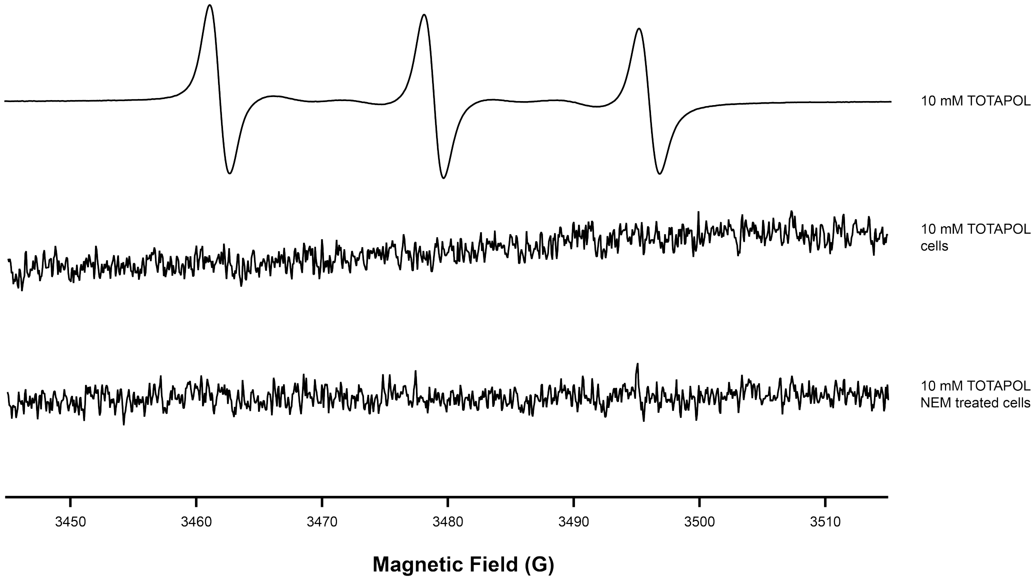


**Figure S7:** EPR spectra showing that TOTAPOL is not taken up into cells prepared identically to DNP samples using CrOx. CrOx is excluded from the cell, broadening TOTAPOL signals that originate from outside, but not inside of the cell. The bottom two EPR spectra do not show TOTAPOL peaks (as compared to TOTAPOL in buffer (top)), indicating that TOTAPOL was not taken up into the cytoplasm in significant concentrations on the timescale used for DNP sample concentration.

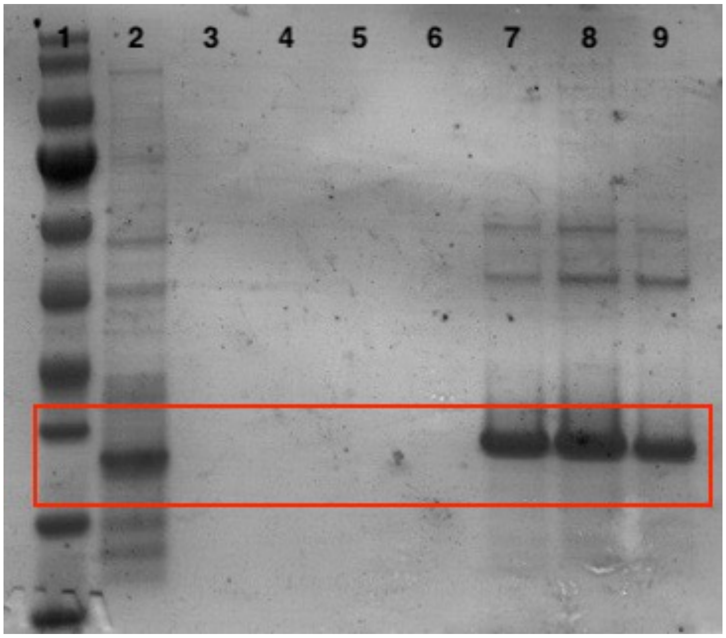


**Figure S8:** SDS-PAGE gel showing significant overexpressed DHFR leakage into cellular media upon treatment with 10% DMSO. Lanes are as follows: 1) *rec* protein ladder 2) cell pellet pre wash 3) NEM treated cells wash 1 supernatant 4) untreated cells wash 1 supernatant 5) NEM treated cells wash 2 supernatant 6) untreated cells wash 2 supernatant 7) NEM treated cells + 10 % DMSO supernatant 8) NEM treated cells + 10 % DMSO supernatant (second centrifugation) 9) untreated cells + 10% DMSO supernatant
